# Diagnosis of coronary heart diseases using gene expression profiling; stable coronary artery disease, cardiac ischemia with and without myocardial necrosis

**DOI:** 10.1101/019505

**Authors:** Nabila Kazmi, Tom Gaunt

## Abstract

Cardiovascular disease including coronary artery disease and myocardial infarction is one of the leading causes of death in Europe, and is influenced by both environmental and genetic factors. With the advancements in genomic tools and technologies there is potential to predict and diagnose heart disease using molecular data from analysis of blood cells. We analyzed gene expression data from blood samples taken from normal people (n=21), non-significant coronary artery disease (n=93), patients with unstable angina (n=16), stable coronary artery disease (n=14) and myocardial infarction (MI; n=207). We used a feature selection approach to identify a set of gene expression variables which successfully differentiate different cardiovascular diseases. The initial features were discovered by fitting a linear model for each probe set across all arrays of normal individuals and patients with myocardial infarction. Three different feature optimisation algorithms were devised which identified two most discriminating sets of genes one using MI and normal controls (total genes=8) and another one using MI and unstable angina patients (total genes=17). The results proved the diagnostic robustness of the final feature sets in discriminating not only patients with myocardial infraction from healthy controls but also from patients with clinical symptoms of cardiac ischemia with myocardial necrosis and stable coronary artery disease despite the influence of batch effects and different microarray gene chips and platforms. selection approach to identify a set of gene expression variables which successfully differentiate different cardiovascular diseases. The initial features were discovered by fitting a linear model for each probe set across all arrays of normal individuals and patients with myocardial infarction. Three different feature optimisation algorithms were devised which identified two most discriminating sets of genes one using MI and normal controls (total genes=8) and another one using MI and unstable angina patients (total genes=17). The results proved the diagnostic robustness of the final feature sets in discriminating not only patients with myocardial infraction from healthy controls but also from patients with clinical symptoms of cardiac ischemia with myocardial necrosis and stable coronary artery disease despite the influence of batch effects and different microarray gene chips and platforms.

## Introduction

Cardiovascular disease (CVD) annually causes about 17.3 million deaths in worldwide and is known as the leading cause of mortality (Mozaffarian et al. 2015). An acute myocardial infarction (AMI) is a necrosis of myocardial tissue due to reduced blood supply to the heart and causes around 735,000 heart attacks in the US every year (Mozaffarian et al. 2015). A large number of scientific advances have been made to prevent, diagnose and treat AMI but unfortunately it is still a leading cause of worldwide morbidity and mortality.

The current diagnosis of AMI is based on potential clinical symptoms including chest pain and impaired breathing and changes in the pattern of ECG and a significant rise and subsequent fall in the circulating levels of cardiac troponins (cTns) (Thygesen et al. 2007). Despite the advances in the cardiovascular field there are several limitations in the current diagnostic system. The advancements in hs-cTn assays have made it possible to detect 10 fold lower circulating Tn concentrations but it has also elevated the number of cardiovascular patients by counting clinically non-diseased people showing changes in cTns due to other conditions (Eggers et al. 2009). Another diagnostic measure is the detection of cardiac miRNAs, which are introduced as sensitive biomarkers (Wang et al. 2008) but their successful detection is inhibited due to their low abundance, small size and tissue specific expression. Their role can be fully advantageous with the invention of fast standardised and automated detection systems (Planell-Saguer and Rodicio 2013). BNP, CRP and other serum inflammatory markers are also considered as cardiovascular biomarkers but they have just slightly improved the diagnosis (Wilson et al. 2008; Shah et al. 2009; Melander et al. 2009)

The majority of cardiac biomarkers are developed using the knowledge of pathological and physiological processes in established pathways. In contrast microarray platforms measure the expression of a large number of genes simultaneously, enabling gene expression profiling across many pathways in parallel. This approach has the potential to represent a comprehensive range of pathophysiological processes of cardiovascular diseases economically and efficiently (Pedrotty et al. 2012). Gene expression profiling extends beyond known biomarkers to reveal biomarkers which are not already known and not previously associated to heart disease.

Gene expression analysis can help to understand and discover novel and sensitive biomarkers of cardiovascular disease. A gene expression analysis yielded 482 genes with an association to the composition of coronary atherosclerotic plaques and most of them were not previously linked to atherosclerosis (Randi et al. 2013). A wide-scale gene expression profiling identified fifty six divergent genes for atherosclerotic and non-atherosclerotic human coronary arteries, wherein 49 were never associated with CAD before (Archacki et al. 2003). Elashoff and co-authors identified a set of classifying genes which with the information of age and sex were strongly correlated to obstructive CAD in non-diabetic patients (Elashoff et al. 2011). The divergent gene expressions were identified which discriminated ischemic and non-ischemic cardiomyopathies conditions among the end-stage patients (Kittleson et al. 2004; Kittleson et al. 2005). Microarray analysis and gene expression profiling were used to discover genes related to heart failure using the expression profiles of 12 patients with heart failure (Min et al. 2010), another study of normal controls and AMI patients discovered genetic markers and dysregulated pathways associated with disease recurrence in first time AMI patients (Suresh et al. 2014).

Despite a range of studies exploring differential expression in cardiovascular outcomes no attempt to use this information to classify patients according to outcome (eg unstable angina and AMI) has been reported yet. If successful this approach offers the potential to provide a diagnostic tool to sub-classify patients. In this work, we identified the discriminatory features to differentiate among normal controls, patients with AMI, stable coronary artery disease (CAD) and unstable angina using gene expression in blood cells. This paper also discusses the success and classification accuracy of different proposed algorithms implemented to discover the potential divergent gene expression features of heart diseases and their optimisation to explore the subset of most discriminatory features.

## Methods-Incorporated Datasets

**“Nelson Dataset”** has gene expression data from blood samples taken from 47 subjects including 26 first time AMI patients and 21 normal people (Suresh et al. 2014). The dataset was subdivided into 2 datasets NelsonA (n=30; 15 first time AMI and 15 controls) and NelsonB (n=17; 11 first time AMI and 6 controls). NelsonA was used as a training dataset to build the classifier and NelsonB was used to identify the final classifier and for the independent validation. The accession number of the dataset at NCBI Gene Expression Omnibus (GEO) is GSE48060. **“Rothman Dataset”** The mRNA expression levels from whole blood of 26 patients with acute coronary syndromes (ACS) were taken at 7 and 30 days post ACS. The dataset has in total 52 samples including eight patients of unstable angina and eighteen of myocardial infarction (MI) (GEO; GSE29111). The samples of both data sets Nelson and Rothman were processed on Affymetrix Human Genome U133 plus 2.0 platform and were normalised using the robust multi-array averaging (RMA) method of affy package in R (Irizarry et al. 2004). **“Beata Dataset”** The blood samples were collected from twenty eight patients of ST-segment elevation myocardial infarction (STEMI) and from fourteen controls which were patients of stable coronary artery disease (CAD). The samples of STEMI patients were taken on first day, after four-six days and after six months of their admission giving a total of 98 samples were processed on Affymetrix GeneChip Human Gene 1.0 ST (GEO; GSE62646). **“Gregg Dataset”** A research group collected blood samples of 338 subjects with mixed cardiovascular phenotypes including non-significant CAD (FINE; n=93), significant CAD, AMI (n=61) and old MI. RNA expression levels were measured on Illumina HumanHT-12 V4.0 expression beadchip (GEO; GSE49925). In our study, we only used FINE and AMI patients (n=154) for independent validation of the classifier.

**Classifier Development-**The expression measures of NelsonA were used to extract the initial differential features and to build the classifier. We used NelsonA to optimise the initial set of features in three different optimisation techniques and generated several subsets. We then used Nelson (NelsonA as a training set and NelsonB a test set) and Rothman (n=24, training set and n=28 test set) to select the final features, the Subset 1 and the Subset 2 based on their highest classification accuracy. The Subset 1 was independently validated on Rothman, Beata, Gregg datasets and the Subset 2 on NelsonB and Gregg in two approaches including blind validation and leave one out cross validation (LOOCV). In our all classification approaches, we applied a k-nearest neighbour (KNN) classification method (kept number of neighbours = 3) (Ripley 1996; Venables and Ripley 2002). A flow chart explaining the discovery and validation process of these two classifiers is given in Figure 1 and a detailed description of the methodology is written below.

**Figure 1:**
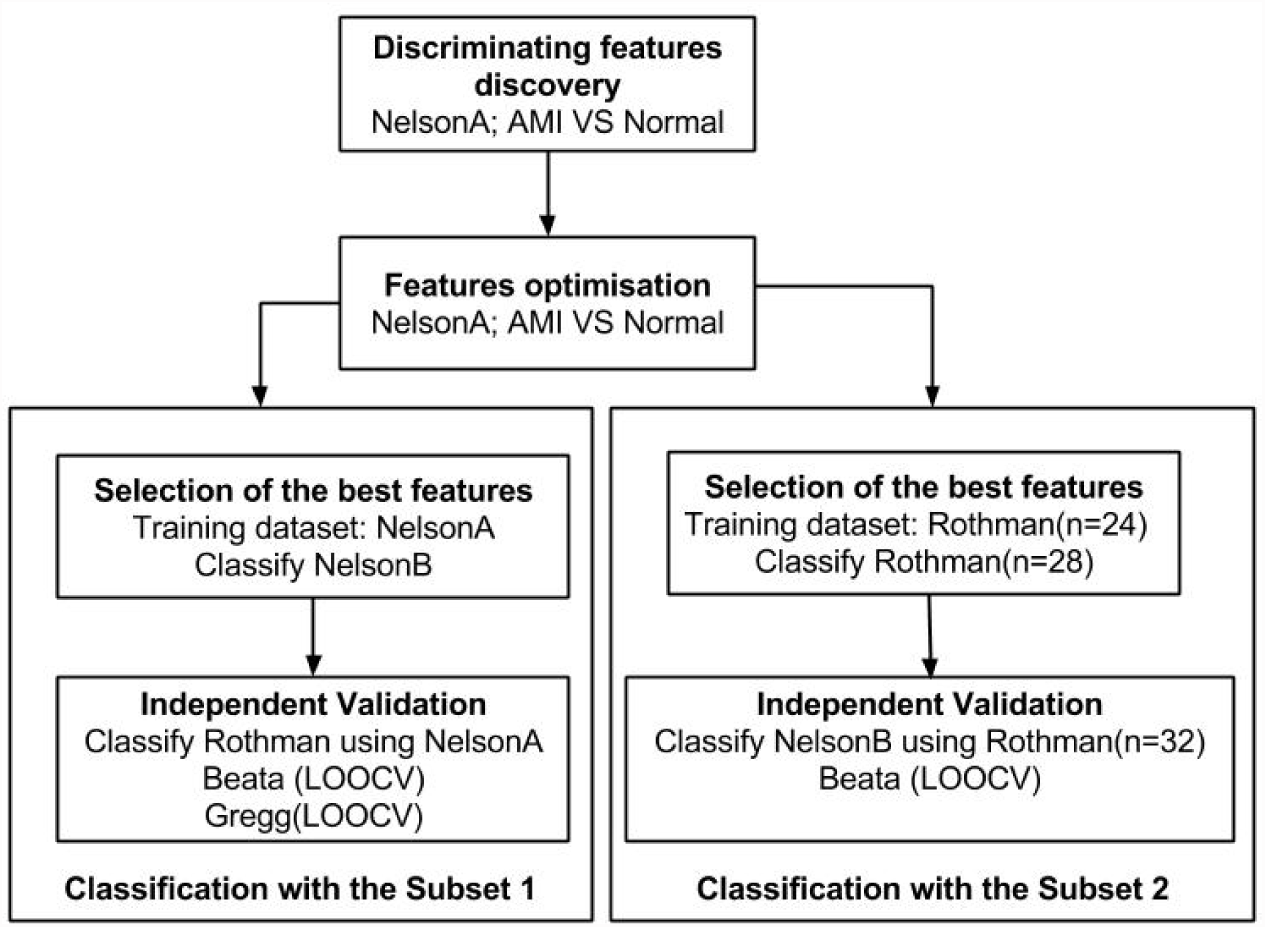
A flow chart to describe the classification process.

**Features Selection (Discovery Phase)-**Initial divergent features were discovered using two different approaches which are described below.

1. **P-Values based Selection (Discovery Method 1)-**A linear model (lmFit) was fitted to training samples and then an empirical Bayes method was used to compute the p-values corresponding to the t-statistics of differential expression (Smyth 2005). The probe sets (PS) were ranked according to their most significant p-values, which were adjusted by using Benjamini and Hochberg (BH) method. The BH method controls the expected false discovery rate (FDR) below a specified threshold and is considered an appropriate choice for microarray experiments (Benjamini and Hochberg 1995). The top 600 genes were selected as the initial classifier.
2. **LOOCV t-tests (Discovery Method 2)-**A linear model (lmFit) was fitted in a LOOCV manner to all training samples (Willenbrock et al. 2004)a. Features were ranked based on the most significant BH adjusted p-values and the most discriminatory genes were retrieved by their maximum number of appearances in the top 300 genes of LOOCV t-tests. This method gave 477 genes. **Features Optimisation-**We developed three different optimisation algorithms to identify the most discriminatory features in two basic lists identified in Discovery Method 1 and Discovery Method 2. All methods processed both lists individually. NelsonA dataset was used to optimise the lists.

1. **Optimisation on Success Rate [Optimisation 1]-**Initiated with the top two PS and tested using LOOCV on the NelsonA samples. Then on the next iteration, the next PS was included and tested using LOOCV again. This was repeated until all PS were processed. After processing of each new PS, we recorded the classifiers success rate (SR; classification accuracy). PS which impaired the performance were eliminated from the list and were given no opportunity for any further LOOCV. The optimisation of features discovered in Method 1 reduced 600 PS to 155 and the features of Method 2 to 123PS.

~~~
nitialisation;
~~~

~~~
currentGene ← 2
~~~

~~~
geneList ← classifying probesets
~~~

~~~
**while** currentGene ≤ totalGenes **do**
~~~

~~~
  Perform *LOOCV* using 1,…,currentGene
~~~

~~~
  Record SR
~~~

~~~
  **if** current SR ≤ previous SR **then**
~~~

~~~
    Eliminate currentGene from the geneList
~~~

~~~
  **endif**
~~~

~~~
  currentGene ← currentGene + 1
~~~

~~~
  **endwhile**
~~~
2. **Performance Improvement and Random Crossover [Optimisation 2]-**The objective of this and the next optimisation technique was to find an optimised set of 25 PS. The method divided all PS into fixed size subsets. For each subset, LOOCV was performed with 1-2,…, n PS of the subset and then the SR and performance index (PI; the performance difference of current and previous PS) was calculated on each iteration. The PS, which impaired the performance was eliminated from the list used for the cross validations but it kept its place in the original subset for evaluation after the crossovers. After processing of the subset, all PS were ranked according to their performance index and the top 25 were retained to perform LOOCV and recorded as an optimised subset on showing a satisfactory SR. The process was evolved for a predefined number of iterations to discover the best combinations. **Crossover**: 2 point crossover was performed between adjacent subsets. After all crossovers the new subsets were evaluated using the cross validation.

~~~
Initialisation;
~~~

~~~
currentGene ← 2
~~~

~~~
Create initial population
~~~

~~~
**repeat**
~~~

~~~
**while** current subset ≤ total subsets **do**
~~~

~~~
Temporary ← current subset
~~~

~~~
**while** currentGene ≤ size of the subset **do**
~~~

~~~
Perform *LOOCV* using 1,…,currentGene of Temporary
~~~

~~~
Record SR and PI
~~~

~~~
**if** current SR ≤ Previous SR **do**
~~~

~~~
Eliminate currentGene from Temporary
~~~

~~~
**Endif**
~~~

~~~
currentGene ← currentGene + 1
~~~

~~~
**endwhile**
~~~

~~~
Rank genes of Temporary according to PI
~~~

~~~
Perform *LOOCV* using top 25 genes of Temporary
~~~

~~~
**If** SR of 25 genes ≥ acceptable **do**
~~~

~~~
Retain the subset
~~~

~~~
**endif**
~~~

~~~
**endwhile**
~~~

~~~
Perform 2 point *crossover*
~~~

~~~
Create new population
~~~

~~~
**until a** predefined number of iterations
~~~
3. **Performance Improvement and Crossover of the Fittest [Optimisation3]-**This method works in a similar pattern as Optimisation 2 and differs only in the crossover technique. **Crossover**: Each subset ranked its PS according to the performance index and performed crossover of its PS with best performing PS of other subsets. After completing all crossovers the newly evolved subsets were evaluated using cross validation. Initialisation;

~~~
currentGene ← 2
~~~

~~~
Create initial population
~~~

~~~
**repeat**
~~~

~~~
    **while** current subset ≤ total subsets **do**
~~~

~~~
     Temporary ← current subset
~~~

~~~
     **while** currentGene ≤ size of the subset **do**
~~~

~~~
       Perform *LOOCV* using 1,…,currentGene of Temporary
~~~

~~~
       Record SR and PI
~~~

~~~
       **if** current SR ≤ Previous SR **do**
~~~

~~~
         Eliminate currentGene from Temporary
~~~

~~~
       **endif**
~~~

~~~
       currentGene ← currentGene + 1
~~~

~~~
       **endwhile**
~~~

~~~
       Rank genes of Temporary according to PI
~~~

~~~
       Perform *LOOCV* using top 25 genes of Temporary
~~~

~~~
       **If** SR of 25 genes ≥ acceptable **do**
~~~

~~~
         Retain the subset
~~~

~~~
          **endif endwhile**
~~~

~~~
       **while** subset < total subsets **do**
~~~

~~~
        Rank genes of all subsets according to PI
~~~

~~~
        Perform crossover of top genes
~~~

~~~
       **endwhile**
~~~

~~~
       Create new population
~~~

~~~
       **until** a predefined number of iterations
~~~

## Results-Selection of the Final Classifiers

The subjects of NelsonA were used as a standard training set to classify NelsonB in order to identify the final classifier among all optimised lists. We identified a set of eight PS which were originally identified using Optimisation 3 on the features of Discovery Method 1 and resulted in 95% SR and AUC = 0.9. In the paper, we called the list **Subset 1**. The Subset 1 classified all AMI patients correctly with 1.0 sensitivity. Another subset of 14 PS (**the Subset 2**; Optimisation 3 and Features Discovery Method 1) was identified among the optimised lists after the classification of Rothman. The Subset 2 classified MI patients from patients with unstable angina with the highest SR = 88% and AUC was 0.79. Random sampling was performed for 15 times and 24 samples of Rothman dataset were selected as a training set which classified an independent set of Rothman (n=28) samples. Random sampling was performed to ensure that the achieved performance was not due to the sampling effects.

A performance comparison of several optimised lists is given in Table 1, these measures were used to select the final classifiers. The enrichment analysis of the genes identified in Subset 1 and Subset 2 was done using a web tool BioMart and the enrichments network is shown in Figure 2. We compared the classifying PS lists with 157 probe sets identified in Nelson work (Suresh et al. 2014) and no overlap was seen.

**Table 1.** The SR of the optimised feature lists on Nelson and Rothman data sets.

| Dataset | Opt1D1 | Opt1D2 | Opt2D1 | Opt2D2 | Opt3D1 | Opt3D2 |
| --- | --- | --- | --- | --- | --- | --- |
| Nelson | 59% | 71% | 89% | 89% | **95%** | 89% |
| Rothman | 75% | 73% | 85% | 84% | **88%** | 85% |

**Figure 2:**
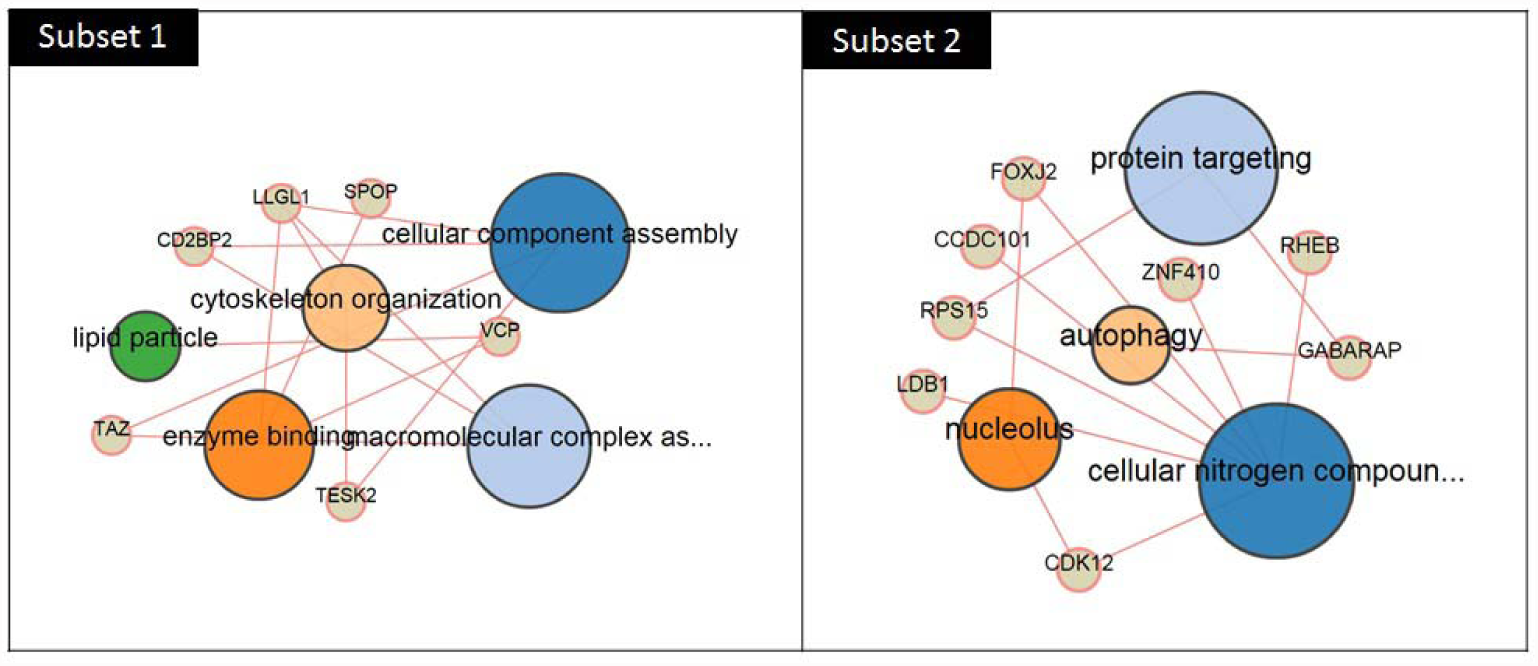
The enrichment network of Subset 1 and Subset 2.

## Independent Validation

The Subset 1 was used to classify the independent data set Rothman (n=52) using NelsonA dataset as a standard training set (Blind Validation). The batch effects were adjusted using COMBAT, an Empirical Bayes method (Johnson and Li 2007). The Subset 1 resulted in 62% SR, sensitivity = 0.75, False Positive Rate (FPR) = 0.69 and precision = 0.71. Then the Subset 2 was used to classify NelsonB using the expression measures of almost 2/3 of Rothman data set (n=32) as a training set. The SR, sensitivity, FPR and precision were 65%, 0.82, 0.67 and 0.7 respectively.

For the classification of Beata dataset (n=98), we mapped our classifying probe sets to HuGene-1_0-st platform using BioMart (http://www.biomart.org/). We successfully mapped all probe sets of the Subset 1 but three probe sets of the Subset 2 were missing. The gene expressions were normalised using RMA method. We called CAD patients our controls and STEMI the cases, performed 15 times random sampling and classified all samples in a LOOCV manner. The random sampling was included to keep the groups balanced for the KNN. The average classification accuracy for the Subset 1 and the Subset 2 was 88% and 96% respectively.

For the classification of Gregg data set, all classifying probe sets of the Subset 1 were mapped to Illumina platform using BioMart. The average bead signals were transformed to log2 scale and were normalised using quantile normalisation (Yang and Thorne 2003). 14,343 probes among 47,211 were constantly detected above the background. The subjects were recruited in two phases and 14,111 probe sets were common in both phases. Unfortunately, one of our mapped probe set was among these 232 missing probes. We considered individuals with FINE phenotype as our controls and AMI as the cases. The LOOCV was performed using the Subset 1 and classified 154 patients after performing fifteen time random sampling. The recorded average SR was 65%.

The receiver operating characteristics (ROC) curve was plotted to measure the quality of both classifiers (Figure 3). The ROC analysis categorised MI patients as positive case and non-MI as negative cases.

**Figure 3:**
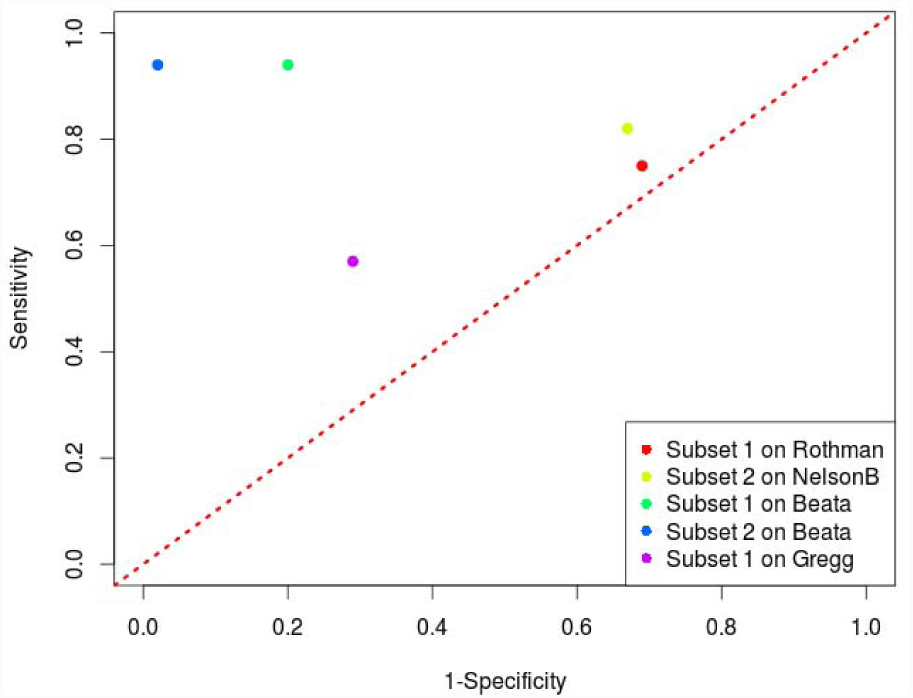
The values of sensitivity and 1-specificity of both classifiers were plotted to evaluate their quality and robustness.

### Discussion

We used NelsonA data set to discover the divergent features between normal controls and AMI patients using two different methods; Discovery Method1 and Discovery Method 2. We identified two separate classifying lists where 461 probe sets were overlapping but were at different positions. We then applied three different optimisation techniques on both lists using NelsonA dataset and generated six different optimised lists. Then, we used Nelson and Rothman datasets to select our final classifiers and the selection criteria was the highest classification accuracy. The selected classifiers were independently validated on Nelson, Rothman, Beata and Gregg datasets following two procedures LOOCV and blind validation.

The ROC analysis was carried out to prove the diagnostic potential of our binary classifiers. In independent validation, the Subset 1 classified Rothman (n=52) using the expression measures of NelsonA (Subset1 on Rothman; Figure 3). The Rothman dataset is a population of patients with unstable angina and MI and we used the expression measures of NelsonA (for training) which has normal controls as negative cases and MI patients as positive cases. The FPR was quite high as expected which indicates our classifier discriminated patients with unstable angina from normal controls of the training set and classified them as MI because their gene expression profiles were closer to positive cases than normal controls. High sensitivity and precision values indicate that the classifier has a high measure of completeness and exactness and is quite robust in predicting AMI from unstable angina with 0.75 recall even under the influence of batch effects.

The Subset 2 classified NelsonB using the expression measures of Rothman (n=32). Using the expression measures of unstable angina (negative cases) and MI patients (positive cases), Subset 2 discriminated positive cases (AMI patients; NeslonB) very well with 0.82 sensitivity (Subset2 on NelsonB; Figure 3). The FPR is again very high because the negative cases of the test dataset (normal controls; NelsonB) were not truly negative. Patients with unstable angina and AMI share many characteristics and our classifier showed a potential to differentiate patients of cardiac ischemia with myocardial necrosis from patients of cardiac ischemia but without myocardial necrosis.

Beata data has a population of STEMI and CAD patients and was classified in a LOOCV manner, where both the training and test samples were from the same dataset. For its classification, Subset 2, which was selected using the information of two diseases (AMI and unstable angina) showed higher measures of both sensitivity and specificity as compared to Subset 1 (Subset1 on Beata, Subset2 on Beata; Figure 3). The Gregg dataset has FINE and AMI patients therefore we considered only Subset 1 for its classification because this list was discovered using the information of AMI and normal individuals (Subset1 on Gregg; Figure 3). The ROC analysis indicates good sensitivity and specificity, indicating its reliability and robustness despite the fact that not all classifying probes sets of Subset 1 could be used and the samples were analysed using a completely different platform. Our results support the utilization of the discovered genes and proposed methods in the diagnosis of cardiovascular diseases using blood gene expression, and suggest potential clinical applications of gene expression data as biomarkers in cardiovascular disease.

### Comparison with Random Forest

We compared our classifier’s performance with a well-known machine learning algorithm Random Forest (Breiman 2001). The Random Forest classifier was first trained using all genes of the Affymetrix Human Genome U133 plus 2.0 chip and the training data set was NelsonA and then on top 600 significant p-valued genes. The cross validation of random forest classifiers on NelsonB showed 59% SR which is substantially weaker than our classifier’s performances (95% SR of Subset 1 on NelsonB using NelsonA as a training set).

This table shows the classification performance of six optimised lists (were optimised using NelsonA) on Nelson and Rothman data sets [Selection of the best features; Figure 1]. Opt1D1 indicates reduced features set after the Optimisation Method 1 on Features Discovery Method 1, Opt2 Optimisation Method 2, Opt3 Optimisation Method 3, D2 Features Discovery Method 2.

